# Constitutive expression of *JASMONATE RESISTANT 1* elevates content of several jasmonates and primes *Arabidopsis thaliana* to better withstand drought

**DOI:** 10.1101/2021.08.23.457374

**Authors:** Sakil Mahmud, Chhana Ullah, Annika Kortz, Sabarna Bhattacharyya, Peng Yu, Jonathan Gershenzon, Ute C. Vothknecht

**Affiliations:** Plant Cell Biology, IZMB, University of Bonn, Kirschallee 1, 53115 Bonn, Germany; Department of Biochemistry, Max Planck Institute for Chemical Ecology, Hans-Knoell-Strasse 8, 07745 Jena, Germany; Crop Functional Genomics, INRES, University of Bonn, 53113 Bonn, Germany; Emmy Noether Group Root Functional Biology, INRES, University of Bonn, 53113 Bonn, Germany; Department of Biochemistry and Molecular Biology, Bangladesh Agricultural University, Mymensingh-2202, Bangladesh

**Keywords:** abiotic stress, feedback regulation, growth-defense trade-off, hormone signaling, jasmonic acid, jasmonyl-isoleucine

## Abstract

Jasmonates have a well-documented role in balancing the trade-off between plant growth and defense against biotic stresses. However, the role of jasmonate signaling under abiotic stress is less well studied. Here, we investigated the function of *JASMONATE RESISTANT 1* (*JAR1*) in drought stress in *Arabidopsis thaliana.* JAR1 converts jasmonic acid (JA) to jasmonyl-L-isoleucine (JA-Ile), the major bioactive form of jasmonates. Comparison of a newly generated over-expression line (JAR1-OE) with *jar1-11*, a T-DNA insertion line in the *JAR1* locus, and Col-0 revealed that constitutively increased JA-Ile production results in stunted growth and a delay in flowering. Upon water limitation, JAR1-OE plants retained more water in their leaves, showed reduced wilting and recovered better from drought stress than the wild type. By contrast, *jar1-11* mutant plants were hypersensitive to drought. RNA-seq analysis and hormonal profiling of plants under control and drought stress conditions provided insight into the molecular reprogramming caused by the alteration in JA-Ile content. Especially JAR1-OE plants were affected in many adaptive systems related to drought stress, including stomatal density, stomatal aperture or the formation of reactive oxygen species (ROS). Overall, our data suggest that constitutively increased expression of *JAR1* can prime Arabidopsis towards improved drought tolerance.

**One sentence summary:** Constitutive expression of *JAR1*primes *Arabidopsis thaliana* to better withstand drought stress

## INTRODUCTION

Jasmonic acid (JA) and its multiple derivatives, collectively called jasmonates, are involved in the regulation of plant growth and development as well as biotic and abiotic stress responses (Zander et al., 2020; Wasternack & Hause, 2013; Koo, 2018). JA biosynthesis is initiated by the formation of 13(S)-hydroperoxylinolenic acid (13-HPOT) from plastidal galacto-lipids through different lipoxygenases (13-LOXs), among which LOX2 is the best described enzyme in Arabidopsis leaves (Wasternack & Hause, 2013; Bell et al., 1995). Subsequently, ALLENE OXIDE SYNTHASE (AOS) and ALLENE OXIDE CYCLASES (AOCs) generate 12-oxo-phytodienoic acid (*cis*-OPDA). *cis*-OPDA is immediately transported into the peroxisome and converted into its reduced form by the OPDA REDUCTASE 3 (OPR3). Subsequent steps involving several enzymes of the ß-oxidation pathway lead to (+)-7-*iso*-JA. After transport into the cytosol, JA is further modified or conjugated to at least 12 different derivatives, including jasmonoyl-L-isoleucine (JA-Ile), 12-hydroxy-JA-Ile (OH-JA-Ile), 12-hydroxy-JA (OH-JA), 12-*O-*glucoside (12-*O*-Glc-JA), 12-HSO_4_-JA and JA-methyl ester (MeJA). All these metabolic products of the jasmonate pathway show varying levels of biological activity (Koo 2018; Wasternack & Hause, 2013).

JASMONATE RESISTANT 1 (JAR1), a member of the GH3 family enzymes, holds a key position in jasmonate biosynthesis because it catalyses the formation of JA-Ile from JA (Guranowski et al., 2007; Staswick & Tiryaki, 2004). JA-Ile exerts its function through the formation of a complex with CORONATINE INSENSITIVE 1 (COI1) and various members of the transcriptional repressor JASMONATE ZIM-domain family (JAZ). In the absence of JA-Ile, JAZ together with various co-repressors binds to different transcription factors (TFs). Accumulation of JA-Ile leads to formation of JA-Ile-COI1-JAZ complexes and releases JAZ-mediated suppression of jasmonate responsive genes (Chini et al., 2007; Thines et al., 2007; Katsir et al., 2008; Yan et al., 2009). Jasmonate-dependent TFs can act as activators and repressors and ultimately regulate hundreds of genes. The bHLH protein MYC2 is considered a master regulator of jasmonate signaling (Dombrecht et al., 2007) that affects many JA-Ile mediated responses. Several of the jasmonate-responsive genes, such as vegetative storage proteins (VSP1 and VSP2), have been shown to be regulated by MYC2 (Wasternack & Song, 2017; Devoto & Turner, 2005). MYC2 is also a target of gibberellic acid (GA) and abscisic acid (ABA) signaling and thus acts as a central hub for transcriptional regulation of many genes involved in plant growth and defense (Wasternack & Song, 2017).

The endogenous JA-Ile concentration is very low, especially compared to other jasmonates (de Ollas et al., 2015b; Balfagón et al 2019). JA-Ile content seems to be tightly controlled via different regulatory loops, including potential auto-regulation of jasmonate synthesis (Hickman et al., 2017). Moreover, the catabolic derivatives of JA and JA-Ile might play a role in maintaining jasmonate homeostasis.

Drought is considered one of the major abiotic stresses that negatively affect plant growth and development (Yang et al., 2010). Tolerance mechanisms to drought comprise a wide range of cellular processes including global reprogramming of transcription, post-transcriptional modification of RNA and post-translational modification of proteins, leading to adaptive alteration of metabolism and plant development (Yang et al., 2010). Stress adaptation often relies on the interplay of multiple hormone signaling pathways (Verma et al., 2016) that balance the trade-off between growth and stress protection (Gupta et al., 2020; Yang et al., 2010; Claeys & Inzé, 2013). Drought tolerance mechanisms are closely correlated with ABA signaling, however, interaction with jasmonate signaling occurs both synergistically and antagonistically depending on the plant organ and stimuli (Yang et al., 2019; Daszkowska-Golec & Szarejko, 2013). Exogenous MeJA application can induce drought responsive genes while *vice versa* the exposure to drought induces jasmonate biosynthesis leading to JA-Ile accumulation (Zander et al., 2020; Clauw et al., 2016; de Ollas et al., 2015b; de Ollas et al., 2015a). ROS production is a common reaction to environmental stresses including drought (Noctor et al., 2014). Drought tolerance mechanisms thus include systems to alleviate ROS damage and JA was found to be involved in activating anti-oxidant mechanisms such as regulating the ascorbate-glutathione cycle (Dombrecht et al., 2007; Sasaki-Sekimoto et al., 2005; Xiang & Oliver, 1998; Savchenko et al., 2019). The allocation of metabolic resources to synthesize plant defense compounds is often associated with reduced growth and biomass accumulation. Therefore, plants have evolved various strategies to balance growth and defense trade-offs to maximize their fitness (Guo et al., 2018; Züst & Agrawal, 2017; Claeys & Inzé, 2013; Zhang & Turner, 2008). The role of jasmonates as the prime regulators of such growth-defense trade-offs has been well established in the case of biotic stresses such as herbivory or pathogen infection (Howe et al., 2018; Züst & Agrawal, 2017, Guo et al., 2018; Wasternack, 2017). Whether jasmonates, especially JA-Ile and its derivatives, are also involved in balancing plant growth and drought tolerance remains largely unknown.

The first mutant in the *JAR1* locus, *jar1-1*, was identified by its insensitivity to exogenous MeJA application in root growth assays (Staswick et al., 1992). Since then, JA-Ile has been shown to be involved in various plant processes such as pathogen resistance, responses to wounding and insect herbivory, as well as in crosstalk with other hormones (Staswick et al., 1998; de Ollas et al., 2015b; Suza & Staswick, 2008). So far, most work exploring the role of jasmonates in growth regulation and stress response either used external MeJA application or used signaling mutants downstream of JA-Ile synthesis such as *coi1* (Howe et al., 2018; Züst & Agrawal, 2017, Guo et al., 2018; Wasternack, 2017). In this work, we used Arabidopsis lines with altered *JAR1* expression to change the endogenous JA-Ile content and thus explore the role of JA-Ile in plant development and drought tolerance. We could show that alteration in JA-Ile content affects plant growth even under non-stress conditions. While a reduced JA-Ile content in the *jar1-11* mutant makes these plants more susceptible to progressive drought, constitutively increased JA-Ile content in a *JAR1* over-expression line strongly alleviates the deleterious effects of drought, making these plants less sensible and more likely to recover. In depth analysis of RNA-seq data obtained under control and early drought stress conditions provided insight into the transcriptional reprogramming caused by the alteration in JA-Ile content. Based on our results, the potential connection between JAR1-dependent changes in gene expression and differences in Arabidopsis growth and drought response phenotypes are discussed. Overall, our data suggest that constitutive JA-Ile production by increased *JAR1* expression can prime Arabidopsis to cope with drought stress.

## RESULTS

### *JAR1* expression levels affect Arabidopsis growth and time of bolting

JA-Ile is the central regulator of the jasmonate signaling pathway, and the JAR1 protein holds a key position in jasmonate biosynthesis because it catalyzes the formation of JA-Ile from JA (Guranowski et al., 2007; Staswick & Tiryaki, 2004). To investigate the effect of JA-Ile on plant growth, we used the Arabidopsis TDNA insertion line *jar1-11* (Suza & Staswick, 2008). Moreover, we generated a line expressing YFP-tagged JAR1.1 under control of the 35S promoter (*35S::JAR1.1-YFP*) in the wild type background, which we refer to as JAR1-OE (**Figure 1 and Supplemental Figure S1**). Three different splice variants have been predicted for *JAR1* that vary slightly in their exon-intron structure (Zander et al., 2020). Of these, JAR1.1 was the first to be identified (Staswick et al., 2002). In *jar1-11*, insertion of the TDNA into the third exon of the JAR1.1 splice variant was validated at the genomic level by PCR (**Supplemental Figure S1A and S1B**). RT-qPCR analysis of rosette leaves under normal growth conditions detected very low expression of *JAR1* transcripts in *jar1-11* (**Figure 1A**), confirming that it is a knock-down but not a null mutant for *JAR1*. By contrast, JAR1-OE plants showed constitutively elevated expression of *JAR1* (**Figure 1A**). Fluorescence microscopy and western blot analysis with a GFP antibody further confirmed the presence of high levels of JAR1.1-YFP protein in rosette leaves of the JAR1-OE line (**Supplemental Figure S1C and S1D**).

**Figure 1.**
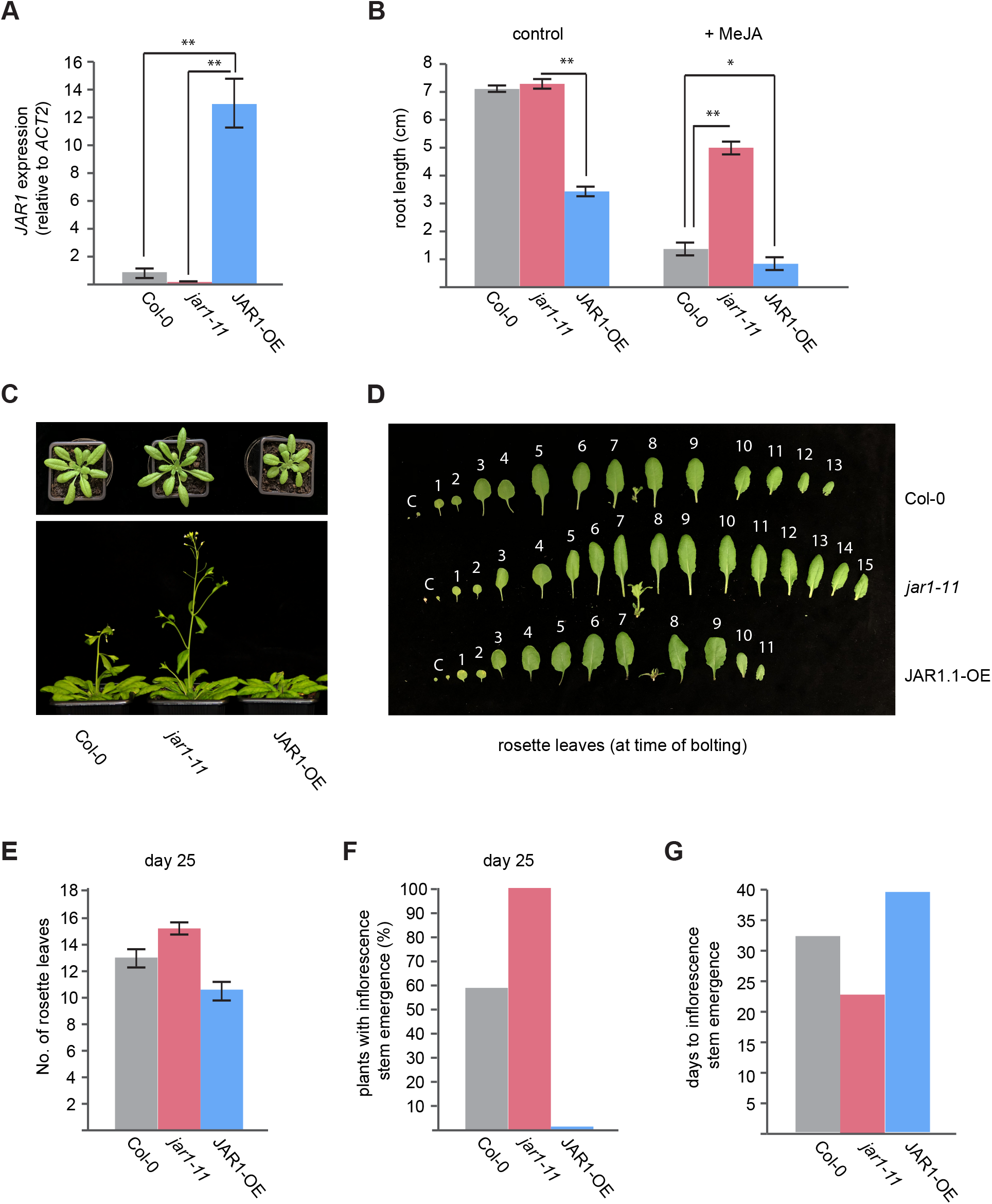
Alteration in *JAR1* expression affects Arabidopsis leaf growth and flowering time. **(A)** *JAR1* transcript levels, relative to *ACT2*, in Col-0, *jar1-11* and JAR1-OE determined by RT-qPCR using rosette leaves of 25 day-old plants grown on soil. Data were analyzed by one-way ANOVA (**P<0.01) followed by multiple comparison analysis (Tukey’s HSD test). Data represent means ±SE from three biological replicates (n=3). **(B)** Root length of Col-0, *jar1-11* and JAR1-OE plants grown on ½ MS medium with or without 50 µM MeJA (see also Supplemental Figure S2). Data were analyzed by one-way ANOVA (*P<0.05, **P<0.01) followed by multiple comparison analysis (Tukey’s HSD test). Data represent means ±SE from three biological replicates (n =3), each containing > 10 seedlings. **(C)** Growth phenotype of Col-0, *jar1-11* and JAR1-OE plants grown on soil under long-day conditions after 25 days (upper panel) and 32 days (lower panel). **(D)** Detached rosette leaves at the time of inflorescence stem emergence (∼ 1 cm length) of plants grown on soil in long-day conditions. Leaves were detached at day 32 (Col-0), 25 (*jar1-11*) and day 40 (JAR1-OE). **(E)** Rosette leaf number of the different plants at day 25. Data represent means ±SE from five biological replicates (n=5), each containing a minimum of five individual plants. **(F)** Percentage of plants with emerged inflorescence stem of at least 1 cm at day 25. **(G)** Average day by which inflorescence stems had emerged. Data represent means from five biological replicates (n=5), each containing a minimum of five individual plants.

We first investigated the growth phenotypes of *jar1-11* and JAR1-OE line compared to the respective ecotype Columbia wild type (Col-0). Several studies on jasmonate biosynthetic and signaling mutants had previously shown a moderate insensitivity towards exogenously applied MeJA in root growth assays (Staswick et al., 1992; Xie et al., 1998). When seedlings were grown on ½ MS medium supplemented with sucrose, *jar1-11* plants grew similar as Col-0, while JAR1-OE plants exhibited a retarded root growth phenotype (**Figure 1B; Supplemental Figure S2)**. Exogenous MeJA application resulted in a strong reduction of root growth and shoot development in Col-0, while the *jar1-11* plants were much less affected and developed quite well. JAR1-OE plants were most severely affected by MeJA treatment (**Figure 1B; Supplemental Figure S2**).

Upon extended growth on soil, *jar1-11* plants displayed a slightly larger rosette size than the Col-0, while JAR1-OE plants showed stunted growth with shorter and somewhat wider leaf blades (**Figure 1C** and **1D**). Moreover, the number of rosette leaves varied, with the highest in *jar1-11* (∼14-16) and the lowest in JAR1-OE (∼10-11) (**Figure 1E**). Also, *jar1-11* plants were a few days ahead in bolting and flowering compared to the Col-0, while JAR1-OE plants lagged behind by about 8-10 days (**Figure 1C, 1F** and **1G**). Even at the time of bolting, JAR1-OE plants still had fewer and shorter leaves compared to both Col-0 and *jar1-11* (**Figure 1D**). We confirmed our observations using the *jar1-12* mutant, a second TDNA insertion line of the *JAR1* locus, and two additional independent YFP-tagged *JAR1.1* overexpression lines (**Supplemental Figure S1A** and **Supplemental Figure S3**). We found similar growth phenotypes, with *jar1-12,* which also had significantly lower *JAR1* transcript levels compared to Col-0 plants (**Supplemental Figure S3A**), growing slightly faster than wild type and showing early flowering, while the JAR1.1 overexpressing lines displaying stunted growth and late flowering (**Supplemental Figure S3B**). All these data indicate that the changes in *JAR1* transcript levels have a strong effect on Arabidopsis growth and development.

### *JAR1* expression levels affect drought tolerance of Arabidopsis

We next used *jar1-11* and JAR1-OE plants to investigate the role of JAR1-mediated JA-Ile formation under drought conditions (**Figure 2** and **Supplemental Figure S4)**. We performed progressive drought experiments by withholding water from 18 day-old well-watered plants grown under 16h/8h long-day conditions (**Figure 2A**). After two weeks of water withholding (day 32) at 40% soil water content (SWC), the first indications of drought effects occurred (**Figure 2B** and **Supplemental Figure S4A**). Hypersensitivity of *jar1-11* to drought became clearly visible at 20% SWC (day 36), with plants displaying stronger signs of wilting compared to Col-0. Three days later, at 10% SWC (day 39), both Co-0 and *jar1-11* plants had reached a state of unrecoverable wilting and re-watering at this stage resulted in 0% survival. By contrast, JAR1-OE plants displayed an extended tolerance to drought and showed signs of wilting only at 10% SWC (day 39). The mild drought effects seen on the JAR1-OE plants at this time point could be fully reversed by re-watering (**Figure 2B** and **Supplemental Figure S4A**). The drought-susceptible phenotype of *jar1-11* could be confirmed in the *jar1-12* line (**Supplemental Figure S4B**). In a separate experiment, the hypersensitivity of *jar1-11* and tolerance of JAR1-OE were also demonstrated by quantifying the leaf relative water content (RWC). At day 36 (20% SWC), Col-0 plants retained around 50% RWC, while the RWC of *jar1-11* plants had dropped to about 30%. By contrast, JAR1-OE plants still remained at 80% RWC (**Figure 2C**).

**Figure 2.**
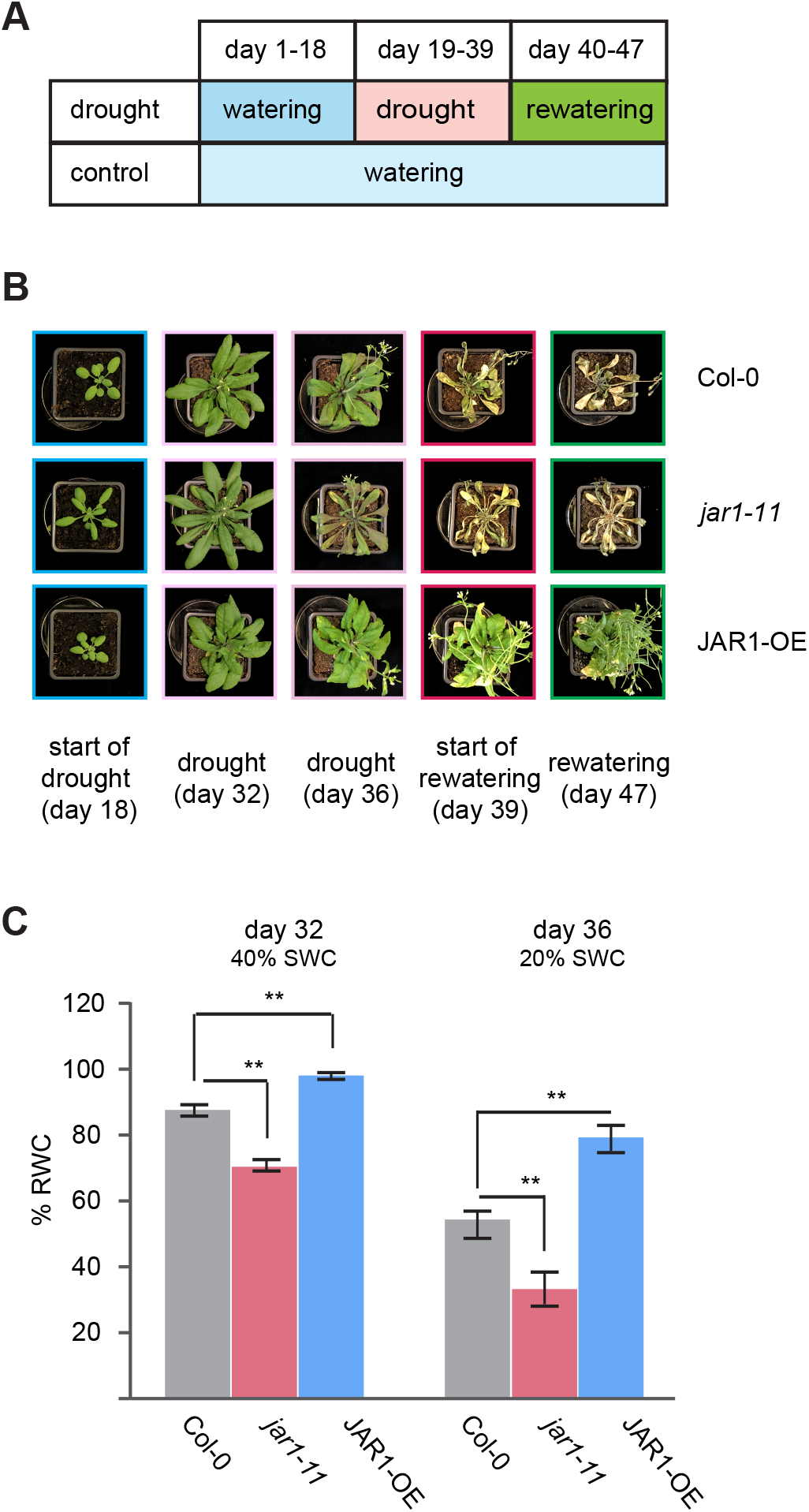
Increased *JAR1* expression positively affects drought stress tolerance. **(A)** Schematic representation of the progressive drought stress experiment. Watering was stopped on day 18. Plants were watered again at day 39 when soil water content (SWC) of Col-0 plants reached 10%. **(B)** Representative photographs showing plant phenotypes throughout the progressive drought stress experiment (see also Supplemental Figure S4A). **(C)** Leaf relative water content (% RWC) of drought-treated plants on day 32 (40% SWC) and day 36 (20% SWC). Data represent means ± SE from five biological replicates (n=5), each containing five individual plants. Data were analyzed by one-way ANOVA (**P<0.01) followed by multiple comparison analysis (Tukey’s HSD test).

We also conducted a similar drought experiment under short-day conditions (**Supplemental Figure S5**). Initially, all lines including JAR1-OE were more heavily affected by water loss than under long-day conditions (**Supplemental Figure S5A**). Already after 14 days of water withholding (day 32), the *jar1-11* plants displayed clear signs of wilting. The Col-0 and JAR1-OE plants showed more tolerance, but all three plant lines were heavily wilted by day 39. None of the *jar1-11* plants recovered after re-watering; however, in contrast to the long-day conditions, most of the Col-0 plants (∼ 80%) were recovered after one week. Again, all the JAR1-OE plants recovered already after 24 h (**Supplemental Figure S5B**).

### JAR1-dependent changes in jasmonates regulate drought tolerance in Arabidopsis

To further elucidate the role of JAR1 in regulating Arabidopsis drought tolerance, we analyzed the contents of various jasmonates in the different plant lines grown under long-day conditions (**Figure 3**). We collected leaf samples of plants grown under control (well-watered) and drought conditions on day 32 (40% SWC), a time point before severe drought symptoms became visible (**Figure 2B**). Under control conditions, JA-Ile in *jar1-11* plants was virtually absent. By contrast, the JAR1-OE plants accumulated elevated levels of JA-Ile, indicating that substantial amounts of JA-Ile were synthesized and retained in the presence of constitutively elevated JAR1.1 protein (**Figure 3D)**. Content of the first committed precursor of jasmonate synthesis, *cis*-OPDA, was slightly increased in JAR1-OE plants compared to Col-0 and the increase was even more pronounced compared to *jar1-11* (**Figure 3G**). On the other hand, content of JA, the direct JAR1 substrate, did not change much, with a slight increase observed in *jar1-11* compared to Col-0 and JAR1-OE (**Figure 3A)**. With regard to JA derivatives, catabolic products such as OH-JA, OH-JA-Ile and COOH-JA-Ile, showed a substantial increase in the JAR1-OE plants (**Figure 3B, 3E and 3F**), suggesting that some of the increased JA-Ile production in these plants led to a greater formation of catabolic products.

**Figure 3.**
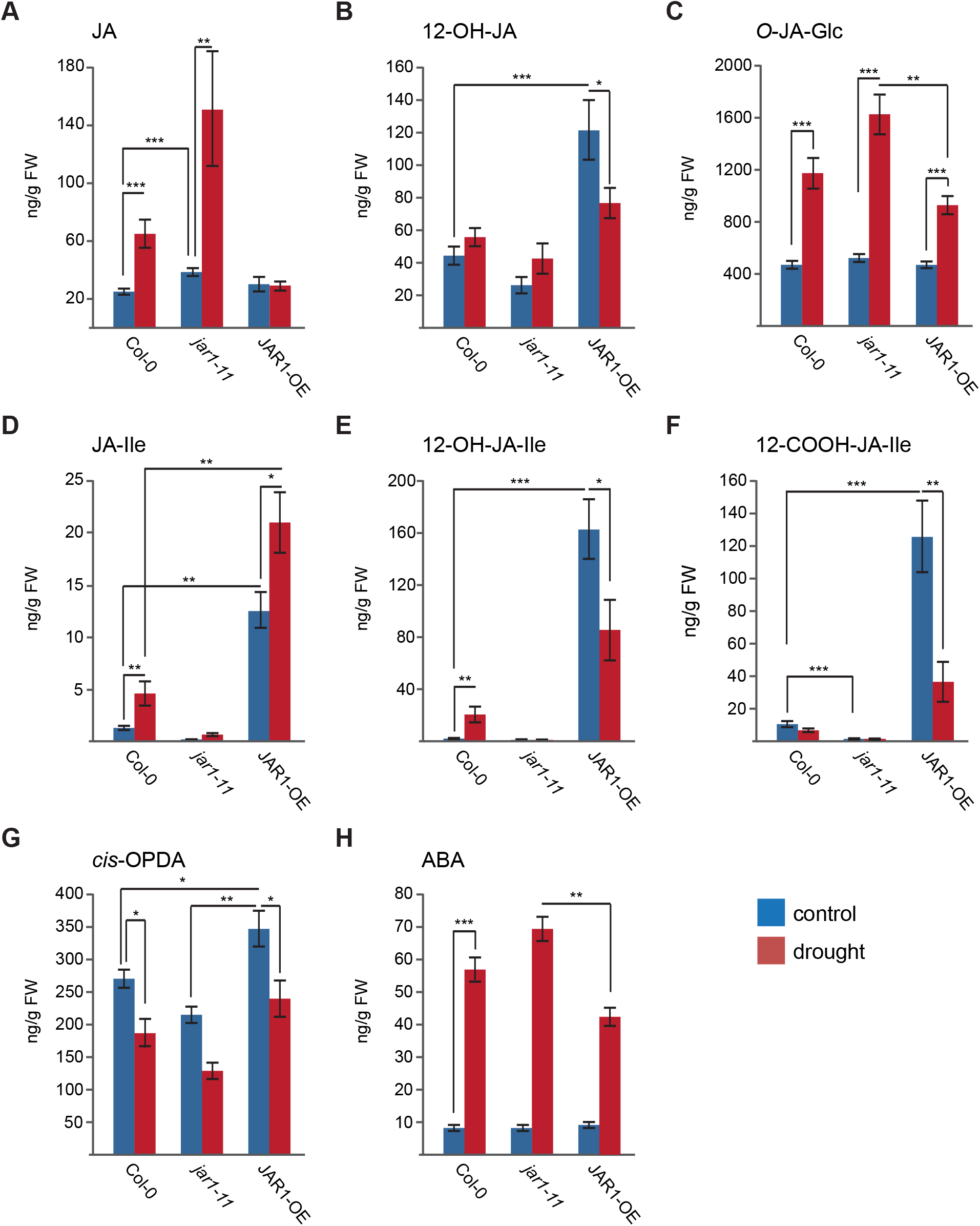
JAR1-dependent changes in the contents of jasmonates and ABA. The contents of different jasmonates **(A-G)** and ABA **(H)** were determined in rosette leaves of 32-day old plants from wild type (Col-0), *jar1-11* and JAR1-OE grown under control and drought stress conditions. Compounds measured were jasmonic acid (JA), 12-hydroxy-jasmonic acid (12-OH-JA), 12-hydroxyl-jasmonyl-glucoside (12-*O*-Glc-JA), jasmonyl-isoleucine (JA-Ile), 12-hydroxy-jasmonyl-isoleucine (12-OH-JA-Ile), 12-carboxy-jasmonyl-isoleucine (12-COOH-JA-Ile), 12-oxo-phytodienoic acid (*cis*-OPDA), and abscisic acid (ABA). Data represent means ± SE from six replicates (n=6), each containing pooled extracts from three plants. Data were analyzed by two-way ANOVA (*P<0.05, **P<0.01, ***P<0.001) followed by multiple comparison analysis (Tukey’s HSD test).

Drought stress resulted in a significant increase in JA-Ile content in Col-0 and JAR1-OE with a proportionally higher increase in Col-0 even though its JA-Ile content under drought was still much lower than in JAR1-OE under control conditions (**Figure 3D**). By contrast, JA-Ile remained virtually absent in *jar1-11* even under drought stress. However, JA content in *jar1-11* was strongly increased since the pathway to JA-Ile is blocked. (**Figure 3A**). JA levels did also increase in Col-0, but not in the JAR1-OE line, likely because increased JAR1 expression allowed more JA to be converted to JA-Ile and its catabolites. Contents of *cis*-OPDA were reduced in all lines under drought (**Figure 3G**) at levels in line with the formation of JA, JA-Ile and derivatives thereof.

*O*-JA-Glc is a less well characterized but highly abundant jasmonates in leaves (Miersch et al., 2008). Its levels were quite similar in all three lines under control conditions and they markedly increased in all lines upon drought stress (**Figure 3C**). Compared to Col-0, the increase was higher in *jar1-11* and lower in JAR1-OE (**Figure 3C**). Similarly, the contents of ABA, which did not differ statistically under control conditions, increased upon exposure to drought stress (**Figure 3H**). Also here the increase was highest in *jar1-11* and lowest in the JAR1-OE plants. Since ABA is the hormone most closely linked to drought stress (Verma et al., 2016), these results suggest that the JAR1-OE line was least affected by drought.

### JAR1-mediated JA-Ile formation regulates many genes involved in Arabidopsis growth and drought tolerance

#### Transcriptional differences between Col-0, jar1-11 and JAR1-OE under normal growth conditions

The strong differences in growth phenotypes of Col-0, *jar1-11* and JAR1-OE plants under normal growth conditions are likely to be controlled by JA-Ile dependent changes in gene expression. To elucidate the global transcriptional changes in these lines, we employed RNA-seq analyses of rosette leaves from 32 day-old well-watered plants grown on soil under long-day conditions (**Figure 4** and **Supplemental Data Set S1**). Using a stringent cut-off (DESeq, adjusted to FDR < 0.01 and LogFC ≥ 1), we found only four differentially expressed genes (DEGs) between *jar1-11* and Col-0 (**Figure 4A; Supplemental Data Set S2**), and all of them were downregulated. By contrast, we found 339 DEGs between JAR1-OE and Col-0 (**Figure 4A, 4B** and **Supplemental Data Set S2**), of which 134 were downregulated and 205 were upregulated. This is in line with the much stronger phenotypic difference observed between JAR1-OE and Col-0 compared to *jar1-11* and Col-0 at this stage under normal growth conditions (**Figure 1C**).

**Figure 4.**
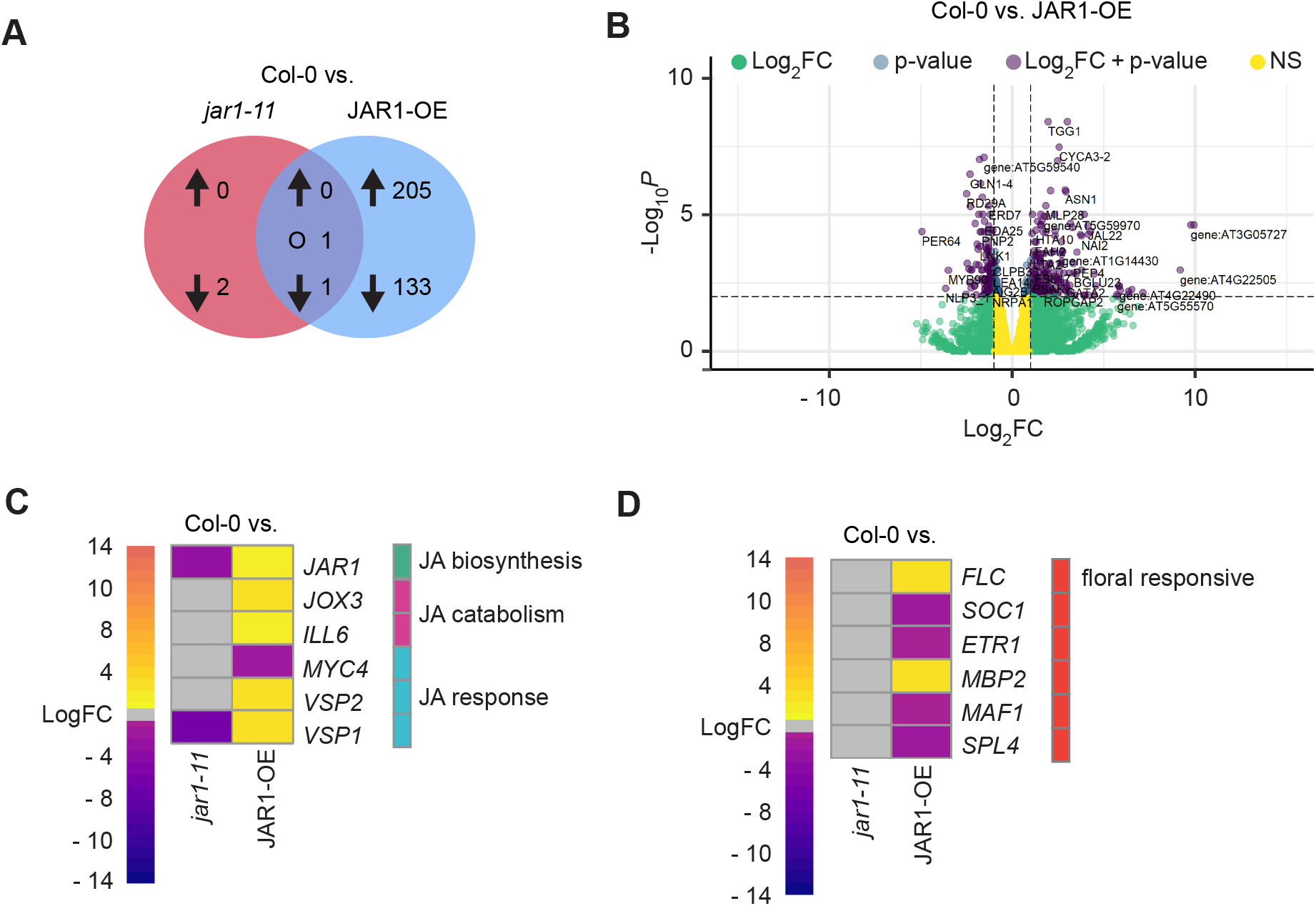
*JAR1*-dependent changes in gene expression in rosette leaves under normal growth conditions. **(A)** Venn diagram showing DEGs (DESeq, adjusted to FDR < 0.01 and LogFC ≥ 1) in *jar1-11* and JAR1-OE compared to Col-0 in 32-day old plants under normal growth conditions. Arrows indicate up- and downregulation. “O” indicates counter-regulated regulated genes. **(B)** Volcano plot showing statistical significance (log_10_*P*) versus magnitude of change (LogFC) of DEGs between Col-0 and JAR1-OE. Violet dots indicate genes that fit the DESeq criteria of FDR < 0.01 and LogFC ≥ 1 while green and blue dots represent DEGs that fit only LogFC or FDR, respectively. **(C) and (D)** Heat maps of genes involved in JA biosynthesis, catabolism and signaling response **(C)** or flowering responsive genes **(D)**. Expression was compared between Col-0 and *jar1-11* or JAR1-OE. Data were analyzed using a cut-off of FDR <0.05 and LogFC ≥ 0.5.

The three genes down-regulated in *jar1-11* (but not JAR1-OE) comprise *JAR1* itself, *AT1G22480* (a potential uclacyanin; cupredoxin superfamily protein) and the well-known jasmonate responsive *VSPI* gene (**Figure 4C; Supplemental Data Set S2**). While the closely related *VSP2* showed only a slight, non-significant decrease in *jar1-11* (**Supplemental Data Set S3**), expression of both, *VSP1* and *VSP2,* was upregulated in JAR1-OE plants (**Figure 4C**). Although it is described that JA-Ile accumulation releases transcriptional repression of *MYC2,* a transcription factor and master regulator of JA-mediated signaling (Lorenzo et al., 2004; Kazan & Manners, 2013), we found only a non-significant increase in *MYC2* expression in the JAR1- OE plants under control conditions (**Supplemental Data Set S3**). Expression of *MYC4*, a transcription factor that was suggested to modulate MYC2-mediated regulation (Fernández-Calvo et al 2011; Wasternack & Hause, 2013), however, was significantly decreased in JAR1-OE (**Figure 4C**). In line with the high levels of JA and JA-Ile derivatives, transcripts of *IAA-LEUCINE RESISTANT (ILR)-LIKE GENE 6* (*ILL6)* and *JASMONATE-INDUCED OXYGENASES 3 (JOX3)* were remarkably higher in JAR1-OE. ILL6, a negative regulator of JA signaling, can hydrolyze JA-Ile and 12-OH-JA-Ile to JA and 12-OH-JA, respectively (Bhosale et al., 2013; Widemann et al., 2013). JOX3 is involved in the oxidation of JA to 12-OH-JA (Smirnova et al., 2017). Interestingly, recent studies have shown that 12-OH-JA and 12-OH-JA-Ile both play a role in the modulation of certain JA-Ile regulated processes (Miersch et al., 2008; Jimenez-Aleman et al., 2019; Poudel et al., 2019).

In line with the observed differences in development and leaf shape, we found that several of the DEGs upregulated in JAR1-OE as compared to Col-0 are involved in cell cycle control (**Supplemental Data Set S2; Supplemental Data Set S3**), for example, *SYP111 (KNOLLE), FBL17, CYCA3;2* and *CYCB1;2* (Gutierrez, 2009). Other genes upregulated in JAR1-OE have functions in cell plate formation (*SYP111* and *CSLD5*), cell wall expansion (*EXPA3)* and cell wall modification *(LTP2)* (Gu et al., 2016; Bernal et al., 2007; Armezzani et al., 2018; Chae et al., 2010). Although *jar1-11* plants showed early and JAR1-OE plants delayed flowering compared to Col-0 (**Figure 1C**), we found no variation in major photoperiod related floral responsive genes such as *FT, LEAFY* or *APETALA2* (Zhai et al., 2015). This might be due to the fact that we analyzed leaf samples. However, some of the genes described as part of the autonomous flowering-time pathway have been shown to be expressed in leaves (Mouradov et al., 2002; Cho et al., 2017) and a heat map using a less stringent cut-off of FDR <0.05 and LogFC ≥ 0.5 shows enhancement of *FLOWERING LOCUS C* (*FLC*) expression in JAR1-OE (**Figure 4D**). Early flowering inhibition by FLC involves repression of *SOC1* (Michaels & Amasino, 2001), whose expression was decreased in JAR1-OE, as was the expression of the early flowering inducers *MAF1* (Ratcliffe et al., 2001) and *SPL4* (Wu & Poethig, 2006). On the other hand, expression of *MYROSINASE BINDING PROTEIN 2 (MBP2; F-ATMBP*), which is related to flowering regulation through the COI1 receptor (Capella et al., 2001), was enhanced (**Figure 4D**). Interestingly, even under control conditions, JAR1-OE plants showed down-regulation of certain drought-(*RD29A, ERD7, LEA14* and *GCR2*) and cold-responsive (*COR15B)* genes compared to Col-0 (**Figure 4B; Supplemental Data Set S3**). These genes have been shown or predicted to be ABA-induced (Mizuno & Yamashino, 2008; Liu et al., 2007; Gaudet et al., 2011) and their down-regulation in JAR1-OE suggests that JAR1 mediated JA-Ile accumulation in the presence of low amounts of ABA (**Figure 3H**) might result in the suppression of some stress response pathways.

#### Transcriptional differences between Col-0, jar1-11 and JAR1-OE under progressive drought conditions

To investigate JAR1-mediated drought tolerance mechanisms, we also conducted RNA-seq analyses using leaf samples of the different Arabidopsis lines exposed to drought stress (**Figure 5** and **Supplemental Data Set S1**). As for hormone analysis, samples were taken on day 32 (40% SWC) before any severe phenotypic effects became visible (**Figure 2B**). In Col-0, we identified 3401 DEGs, of which 2023 were down-and 1378 upregulated between control and drought conditions (**Figure 5A**). By comparison, *jar1-11* plants, which were most heavily affected by drought stress, showed a much higher number (6139 in total; 2616 up- and 3523 down-regulated) of DEGs, while the more drought-tolerant JAR1-OE line displayed a lower number (2025 in total; 971 up- and 1054 down-regulated) of DEGs. Our data indicate that despite any outside appearance of apparent drought effects at the time of sampling, drought had already affected all three plant lines resulting in substantial changes in global gene expression.

**Figure 5.**
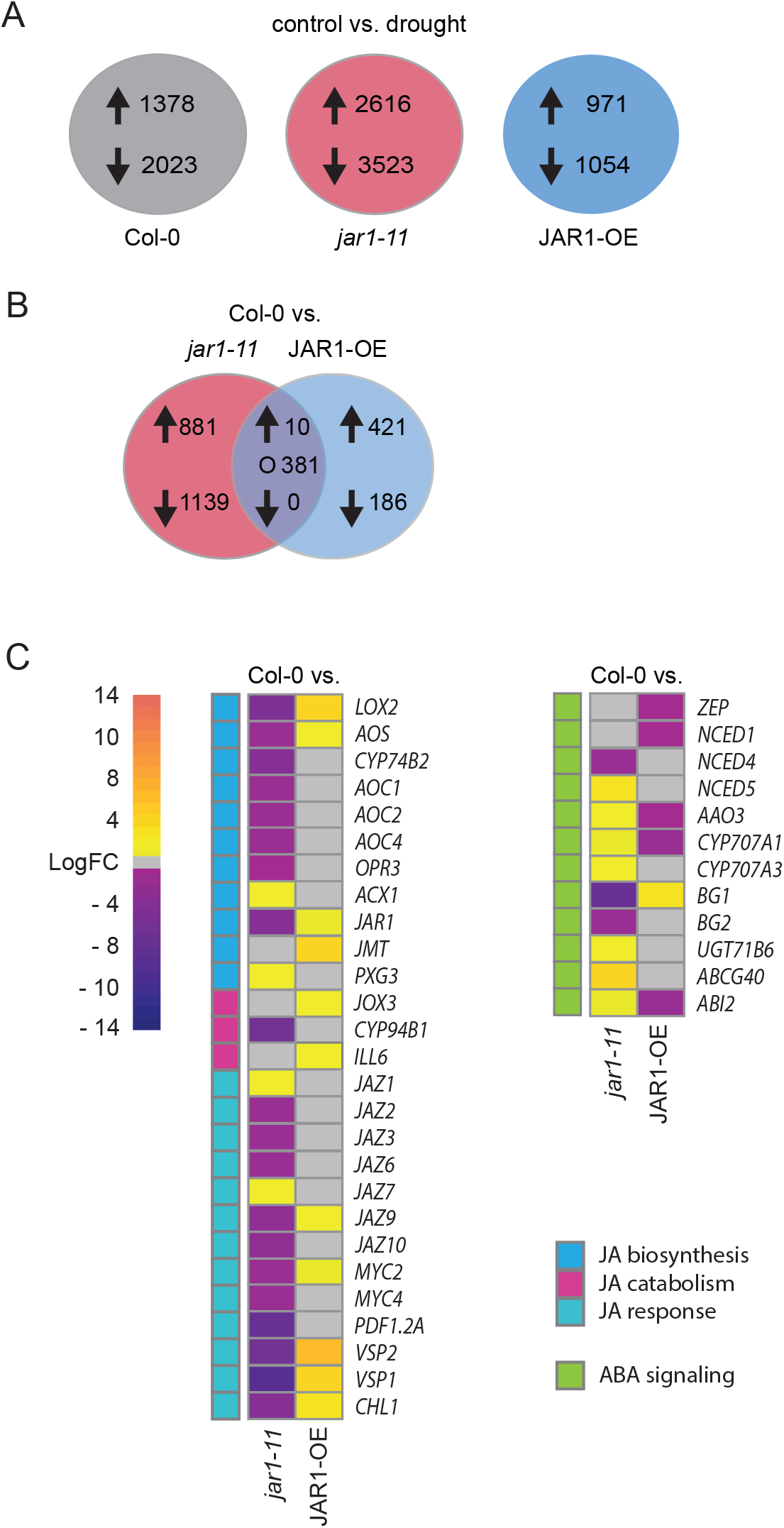
JAR1-dependent changes in gene expression in rosette leaves under progressive drought. **(A)** Number of DEGs (DESeq, adjusted P < 0.01 and LogFC ≥ 1) between control and drought conditions in Col-0, jar1-11 and JAR1-OE. Arrows indicate up- and downregulation. **(B)** Venn diagram of DEGs (DESeq, adjusted P < 0.01 and LogFC ≥ 1) in *jar1-11* and JAR1-OE compared to Col-0 under drought conditions. Arrows indicate up- and downregulation. “O” indicates counter-regulated genes. **(C)** Heat maps of genes involved in JA biosynthesis, catabolism and signaling response (left) as well as ABA biosynthesis, catabolism and signaling response (right) compared between Col-0 and either *jar1-11* or JAR1-OE, all under drought conditions. Data were analyzed using a cut-off of FDR <0.05 and LogFC ≥0.5.

A comparison of the RNA-seq data between the different plant lines under drought conditions revealed 2411 DEGs between wild type and *jar1-11* (**Figure 5B; Supplemental Data Set S2**), among which 966 genes showed a higher and 1445 genes a lower expression level in *jar1-11*. On the other hand, out of 998 DEGs found between Col-0 and JAR1-OE, 737 genes showed a higher and 261 genes a lower expression level in JAR1-OE (**Supplemental Data Set S2**). Moreover, we found 391 DEGs counter-regulated between *jar1-11* and JAR1-OE, most of which showed higher expression in JAR1-OE and lower expression in *jar1-11*. Gene ontology (GO) enrichment analysis confirmed the reciprocal trends between *jar1-11* and JAR1-OE for a number of genes (**Supplemental Data Set S4**), including several genes involved in jasmonate synthesis and jasmonate signaling or known to be jasmonate-responsive (**Figure 5C**). Not surprisingly, expression of the JA responsive transcription factor *MYC2* as well as the MYC2-dependent JA-responsive genes *VSP1* and *VSP2* was upregulated in Col-0 and JAR1-OE but not in *jar1-11* (**Supplemental Data Set S3**). Expression of MYC4, which had been down-regulated in JAR1-OE under control conditions (**Figure 4C**), remained unchanged but was down-regulated in both *jar1-11* and Col-0 (**Supplemental Data Set S3**). In general, differences between Col-0 and *jar1-11* were more pronounced than differences between Col-0 and JAR1-OE, with many jasmonate-related genes showing lower expression in *jar1-11* upon drought (**Figure 5C**). This includes several of the jasmonate biosynthetic genes upstream of *JAR1,* which showed a lower expression in *jar1-11* under drought compared to Col-0, while their expression was similar or higher than Col-0 in JAR1-OE plants (**Figure 5C**). A similar pattern was also observed for the expression of *MYC2, VSP1* and *VSP2* as well as for most JAZ genes. By contrast, several enzymes involved in the formation of JA and JA-Ile derivatives, including *JASMONIC ACID CARBOXYL METHYLTRANSFERASE* (*JMT*), *JOX3*, and *ILL6* showed higher expression in JAR1-OE but similar expression in Col-0 and *jar1-11*. Only a few genes showed higher expression in *jar1-11*. These included the *PEROXISOMAL ACYL-COENZYME A OXIDASE 1* (*ACX1)* which is involved in both jasmonate biosynthesis and ß-oxidation, as well as *PEROXIGENASE 3* (*PGX3*). Also, two *JAZ* genes, *JAZ1* and *JAZ7,* showed a higher expression level in the *jar1-11* mutant (**Figure 5C**).

Not surprisingly, genes known to be responsive to drought and ABA signaling were enriched in the upregulated gene sets of all three lines upon drought (**Supplemental Data Set S2 and S3**). Compared to Col-0 and JAR1-OE, *jar1-11* plants showed a stronger upregulation of several genes involved in the ABA signaling pathway (**Figure 5C**), which is in line with the higher accumulation of ABA under drought (**Figure 3H**). These included genes involved in ABA synthesis, such as *NCED5* and *AAO3,* or *ABI2*, a target gene of ABA regulation (**Figure 5C**). However, we also found that the expression of some genes involved in ABA homeostasis (Xu et al., 2013), such as *CYP707A1*, *CYP707A3* and *UGT71B6,* were higher in *jar1-11*.

The major genes with decreased expression in *jar1-11* and increased expression in JAR1-OE were related to photosynthesis (**Supplemental Data Set S4**). This included the light-harvesting complex genes *LHCB6*, *LHCB2.4* and *LHCB4.2*, whose expressions were significantly lower in *jar1-11* and higher in JAR1-OE compared to Col-0 (**Supplemental Data Set S2 and Set S3**). On the other hand, the major genes with increased expression in *jar1-11* and decreased expression in JAR1-OE included various groups of genes responding to abiotic stresses and other hormones (**Supplemental Data Set S4**). Compared to Col-0, JAR1-OE showed a lower expression of several drought-responsive genes such as *LEA46*, *LEA18*, *LEA7* and *RAB18*. These genes were upregulated under drought conditions in the Col-0 (**Supplemental Data Set S2 and S3**), supporting the phenotypic evidence that the JAR1-OE plants did experience less severe drought stress after 14 days of water withholding. At the same time, transcripts of drought-responsive genes such as *DREB2A* or *RD20*, the drought and cold-responsive gene *COR47,* putative drought-responsive genes such as *LEA31*, and hypoxia-responsive genes such as *FMO1*, *At2g25735* and *HIGD2* had a higher expression level in *jar1-11* compared to Col-0, underpinning the greater susceptibility of *jar1-11* to drought stress.

To further investigate the differential expression in response to drought compared to control conditions, especially between *jar1-11* and JAR1-OE, we applied hierarchical clustering to all DEGs among Col-0, *jar1-11* and JAR1-OE (**Supplemental Data Set S5**). Using the K-means approach, genes were assigned to 5 clusters. The clusters were then visualized with a heat map (**Figure 6**), revealing general patterns of transcriptomic profiles during drought stress compared to control conditions. These clusters can be categorized into two sets, with the first set of clusters (1-4) representing mechanisms to withstand drought stress effects. We found a decreased expression after drought stress in all lines in clusters 1-4, albeit to a lesser extent in JAR1-OE compared to the wild type and especially to *jar1-11*. Many genes in clusters 1 relate to water transport, while cluster 2 and 4 clearly represent the detrimental effect of drought on the photosynthetic machinery. Genes related to growth regulation were affected on several levels from general regulation of growth (cluster 1) to cell wall biosynthesis and remodeling (cluster 3). Cytokinin response was also negatively affected by drought, especially in *jar-11,* supporting a crosstalk between jasmonates and cytokinin under drought stress. Cluster 5 represents drought stress responses and we found upregulation of ABA-dependent and independent genes related to water deprivation (but also genes related to salt stress, hypoxia, heat stress and oxidative stress as well as some jasmonate mediated biotic stresses) in all lines with the highest upregulation in *jar1-11*.

**Figure 6.**
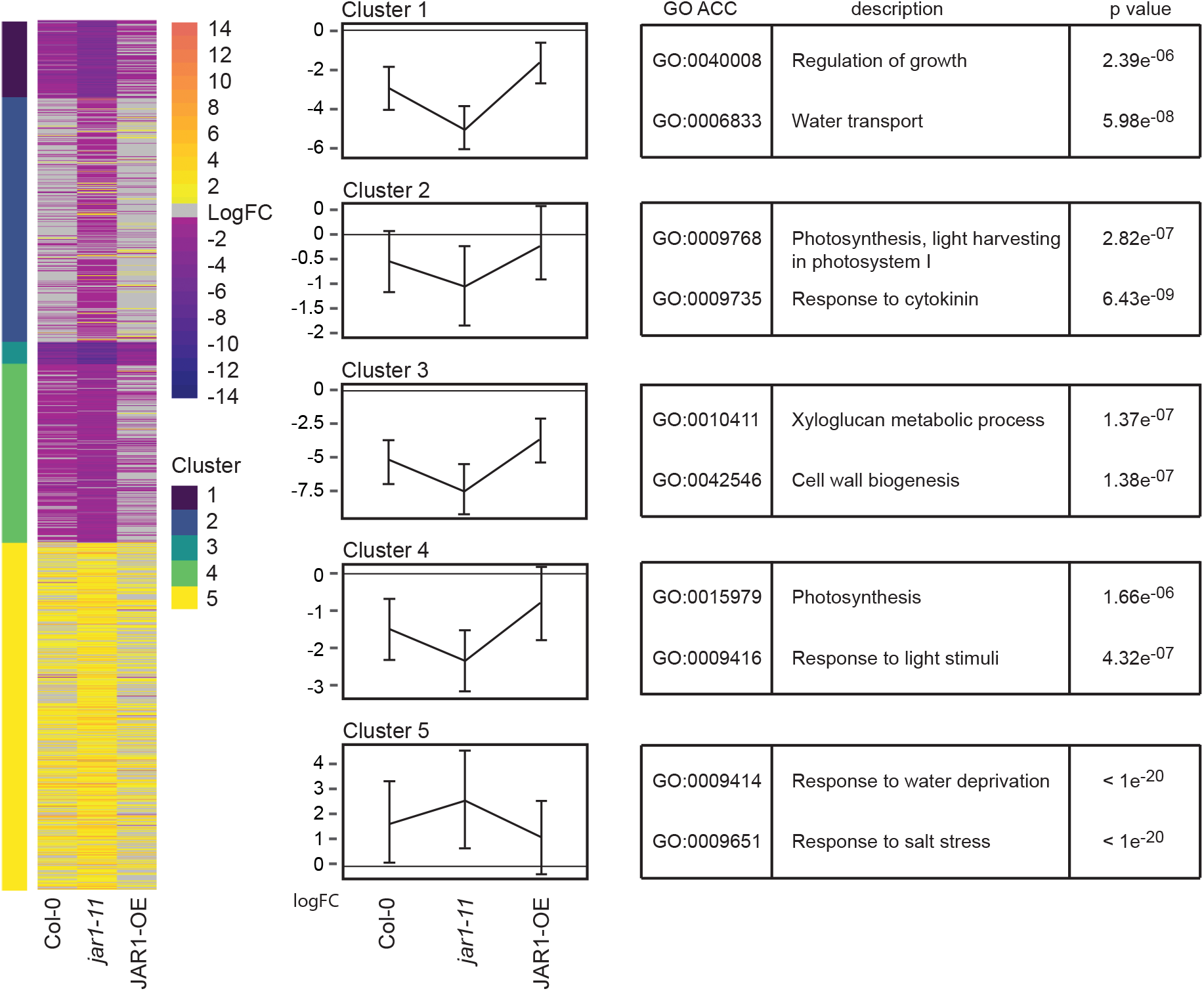
*JAR1*-dependent transcriptomic variations between drought stress and control conditions. **(A)** Heat map (left) and K-means clustering (middle) of genes up- or downregulated under drought stress compared to control conditions in the different plant genotypes. K-means clustering analysis was performed to produce the clusters (DESeq, adjusted FDR < 0.01 and LogFC ≥ 1) and the thin lines represent the mean expression profiles for each cluster (middle). Only genes that are differentially expressed in at least one of the comparisons were used for the cluster analysis. The top two GO terms for each cluster with P values are listed (right).

### JAR1 regulates stomatal aperture and density

Our RNA-seq analysis had revealed that the expression of *TGG1* and *TGG2,* was highly elevated in the JAR1-OE line under normal growth conditions (**Figure 4B** and **Supplemental Data Set S3**). These myrosinases were shown to be involved in ABA- and MeJA-induced stomatal closure downstream of ROS production (Rhaman et al., 2020; Islam et al., 2009). Expression of these genes decreased in all three lines under drought. However, the relative transcript levels in JAR1-OE under drought were still much higher than in Col-0 under control conditions. Similarly, some genes that negatively control stomatal density and distribution, such as *TMM*, *EPF1* or *SBT1.2,* were expressed at higher levels in JAR1-OE under normal growth and drought conditions (**Supplemental Data Set S3**). We thus measured stomatal aperture and density of the 6th rosette leaf of 21 days old Col-0, *jar1-11* and JAR1-OE plants grown under control conditions (**Figure 7**). We found a higher stomatal density in *jar1-11* compared to Col-0 and JAR1-OE (**Figure 7A**), confirming the role of endogenous JA-Ile content on stomata development. Moreover, *jar1-11* plants also displayed a wider stomatal opening (**Figure 7B**). On the other hand, both the stomatal density and aperture diameter were remarkably lower in JAR1-OE compared to *jar1-11* and Col-0 (**Figure 7A and 7B**). Thus, JAR1-mediated JA-Ile accumulation affects both the aperture and density of stomata, which together can affect the transpirational water loss.

**Figure 7.**
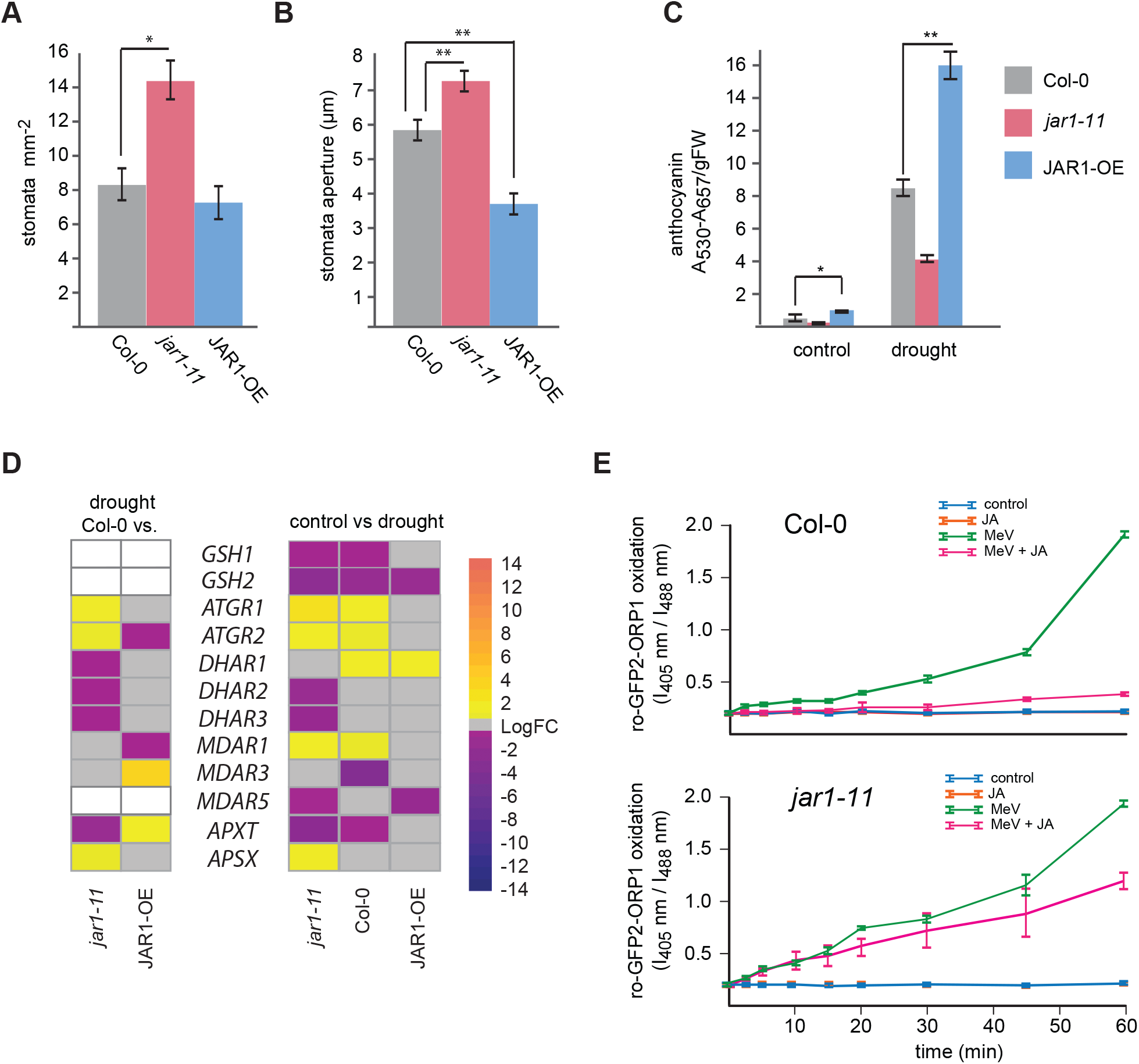
Effect of JA-Ile on stomatal regulation, anthocyanin content and MV-induced changes in redox status. **(A) and (B)** Number of stomata **(A)** and stomatal aperture **(B)** measured on leaf No. 6 of plants grown under control conditions at day 21. Data represent means ± SE from three biological replicates (n=3). For stomatal numbers, each replicate quantified leaves from 5-6 individual plants. For stomatal aperture, each replicate quantified 90 to 100 stomata in leaves from 6-10 individual plants. Data were analyzed by one-way ANOVA (*P<0.05, **P<0.01) followed by multiple comparisons (Tukey’s HSD test). **(C)** Anthocyanin content of different plant genotypes determined in rosette leaves of 32-day old plants grown under control and drought stress conditions. Data represent means ± SE from 3 replicates (n=3), each containing three pooled individual plants. Data were analyzed by one-way ANOVA (*P<0.05, **P<0.01) followed by multiple comparisons (Tukey’s HSD test). **(D)** Heat maps of DEGs involved in the ascorbate-glutathione cycle in *jar1-11* and JAR1-OE compared to Col-0 under drought conditions (left) or between control and drought conditions in Col-0, *jar1-11* and JAR1-OE (right). Data were analyzed using a cut-off of FDR <0.05 and LogFC ≥0.5. White boxes indicate genes whose changes did not meet the cut-off criteria. **(E)** Real-time monitoring of redox status using cytosolic roGFP2-Orp1 redox sensors in Col-0 and *jar1-11* leaf cells upon treatment with 10 mM MeV and/or 1 mM JA. Ratios were calculated from the fluorescence values recorded at 535 nm after excitation at 405 nm and 488 nm. Mean ratios ± SE of different time-points represent data from three replicates, each including three individual seedlings.

### JAR1-dependent modulation of the ascorbate-glutathione cycle

ROS production is a common reaction to environmental stresses, including drought (Noctor et al., 2014). Flavonoids, such as anthocyanins, have been suggested to scavenge ROS, and anthocyanin biosynthesis is induced by MeJA application (Shan et al., 2009). Accordingly, JAR1-OE plants had a higher anthocyanin level under control conditions compared to Col-0 and *jar1-11* (**Figure 7C**). Anthocyanin levels increased significantly in all three plant lines upon drought, with the highest increase observed in JAR1-OE plants. Moreover, glutathione plays an important role in preventing oxidative stress by scavenging H_2_O_2_ through the ascorbate-glutathione cycle pathway (**Supplemental Figure S6**). Some genes coding for enzymes involved in glutathione synthesis or the ascorbate-glutathione cycle pathway were shown to be induced by MeJA application (Xiang & Oliver, 1998; Sasaki-Sekimoto et al., 2005; Zander et al., 2020), while on the other hand, the content of GSH affects MeJA induced stomata closure (Akter et al., 2013). In our experiment, very little difference in expression could be observed between Col-0, *jar1-11* and JAR1-OE under non-stress conditions for genes involved in glutathione biosynthesis or the ascorbate-glutathione cycle (**Supplemental Data Set S3**). Under drought conditions, the GSH producing genes *GSH1* and *GSH2* were downregulated in Col-0 and *jar1-11*, while expression of *DHAR1*, the dehydroascorbate reductase that converts GSH to GSSG, was increased in Col-0 and JAR1-OE (**Figure 7D**). By contrast, expression of *ATGR1* and *ATGR2,* glutathione reductases that convert GSSG to GSH, was increased under drought in Col-0 and to an even greater level in *jar1-11*, but no changes were found in JAR1-OE (**Figure 7D**).

To elucidate possible JAR1-mediated effects on ROS regulation, we used the genetically encoded *in vivo* H_2_O_2_ sensor roGFP-Orp1, which indicates the oxidation level by measuring the H_2_O_2_/H_2_O ratio (Nietzel et al., 2019). We then applied methyl viologen (MV), which was shown to lead to oxidative stress and the generation of ROS, including H_2_O_2_ (Schwarzländer et al 2009). Treatment of leaf tissue from Col-0 plants with 10 mM MV resulted in a strong oxidative shift of the sensor indicative of oxidative stress (**Figure 7E, Col-0, green line**). Application of 1 mM JA, given together with MV, reduced the MV-induced increase in H_2_O_2_ levels nearly back to control levels, indicated by a lack of sensor oxidation (**Figure 7E, Col-0, magenta and blue lines**). Application of JA alone had no effect on H_2_O_2_ levels (**Figure 7E, Col-0, orange line**). Application of 10 mM MV to leaf tissue from *jar1-11* plants carrying the roGFP-ORP1 sensor, resulted in a similar oxidative shift as in Col-0 (**Figure 7E, *jar1-11*, green line**). However, application of JA together with MV only resulted in a minor decrease of sensor oxidation in *jar1-11*, showing that the MV-induced increase in H_2_O_2_ level was not much alleviated (**Figure 7E, *jar1-11*, magenta line**). This indicates that JAR1-mediated transformation of JA to JA-Ile is required to reduce ROS that are induced by MV.

## Discussion

In nature, plants are constantly exposed to various biotic and abiotic stresses. To combat those detrimental effects and maximize their fitness, plants try to strike a balance between growth and stress tolerance mechanisms. Studies had shown that a common set of genes are induced by both externally applied MeJA and drought (Zander et al., 2020; Hickman et al., 2017; Huang et al., 2008). However, the effects of endogenous JA-Ile content on plant growth and drought response remain largely unexplored. Our comparison between *jar1-11* and JAR1-OE demonstrates that JA-Ile plays an important role in regulating processes that help Arabidopsis to withstand progressive drought stress effects. Constitutively increased *JAR1* expression in JAR1-OE plants overrides regulatory circuits that normally reduce JA-Ile content and thus protects plants better from the effects of drought most likely as a result of JA-Ile dependent priming. However, this comes at the cost of retarded growth, delayed flowering and prolonged time until seed production. Therefore, although a consistently high rate of jasmonate signaling due to strong *JAR1* expression might be beneficial for the plant under prolonged drought stress conditions, a more regulated signaling cascade might be a more favorable way to cope with periodic mild drought episodes because it will better promote growth when conditions permit.

### Overexpressing *JAR1.1* induces jasmonate signaling under non-stress condition

In the current work, we used a TDNA insertion mutant in the *JAR1* locus (*jar1-11*) and a novel transgenic line expressing JAR1.1-YFP under the 35S promoter (JAR1-OE) to alter the endogenous JA-Ile content of Arabidopsis. The *jar1-11* mutant showed a strong reduction in *JAR1* transcripts compared to Col-0 (**Figure 1A**), but a basal level of full-length transcripts is retained despite the disruption of the *JAR1* locus within an exon after about 1/3 of the coding region. Some *JAR1* transcript formation had been shown previously for this line under biotic stimuli (Suza and Staswick 2008). Nevertheless, *jar1-11* plants showed a clear reduction in JA-Ile content and nearly null expression of the jasmonate-dependent defense marker *VSP1.* In the JAR1-OE line, strongly increased *JAR1* transcript levels (**Figure 1A**) result in an about 10-fold increase in JA-Ile content (**Figure 3D**), together with upregulation of *MYC2*, *VSP1* and *VSP2*. Thus, these lines are a great resource to study the effects of varying JA-Ile levels on plant growth and stress responses.

### Constitutive alteration in *JAR1*levels alters plant growth and development

The differences in *JAR1* transcript and JA-Ile levels in these transgenic lines manifested themselves in opposite phenotypic alterations compared to Col-0. Differences in growth occurred even under non-stress conditions and were generally more pronounced in JAR1-OE compared to *jar1-11*. Reduced JA-Ile content in the *jar1-11* plants led to a slightly larger rosette size. By contrast, overexpression of *JAR1* resulted in shorter and wider leaves, a similar phenotype also achieved by treating Col-0 plants with exogenous MeJA application (**Supplemental Figure S4C**). Exogenous MeJA application has been shown to arrest the cell cycle and thus growth of young leaves (Noir et al., 2013; Zhang & Turner, 2008). However, the initial stunted growth observed in JAR1-OE seems to be superseded at a later stage by increased radial growth of older leaves. Accordingly, expression of the cell cycle controlling gene *CYCB1.2*, which was found to be downregulated after exogenous MeJA application in young seedlings (Zhang & Turner, 2008), was upregulated in the older leaves of the JAR1-OE plants used for RNA-seq analysis in our experiments. JAR1-OE plants also seem to have higher expression levels of the transcription co-activators *GIF1* and *GRF5_1* (**Supplemental Data Set S3)**, which together were shown to regulate the development of leaf size and shape (Kim & Kende, 2004; Lee et al., 2009). Mutants in the *GIF1* locus had narrower leaf blades compared to wild type indicating that GIF1 supports lateral leaf expansion (Horiguchi et al., 2011; Kim & Kende, 2004; Lee et al., 2009). Altered expression of *GIF1* and *GRF5* in JAR1-OE could be due to the decreased expression of *MYC4*, which was shown to bind the promoter of *GIF1* and downregulate its activity (Liu et al., 2020).

Plants of the *jar1-11* and JAR1-OE lines also display opposite phenotypes with regard to flowering time, which is typically controlled by endogenous factors as well as environmental signals. In our analysis, this difference was not reflected by changes in the expression of known photoperiodic floral inducing genes, even though it had been shown previously that COI1 inhibits *FT* expression (Zhai et al., 2015). However, JA-Ile independent functions of COI1 have been described recently (Ulrich et al., 2021). Instead, the difference in flowering time could be related to vernalization and the autonomous flowering-time pathway, as evidenced by upregulation of the flowering repressor *FLC* and down-regulation of the FLC-repressed transcriptional activator *SOC1* in JAR1-OE (Michaels & Amasino, 2001; Richter et al., 2019).

### JA-Ile regulates several physiological systems involved in drought adaptation and stress response

Clearly, *jar1-11* plants were more susceptible than Col-0 to the progressive drought stress applied in our study, while JAR1-OE plants only displayed a mild drought stress phenotype. A similar drought stress tolerance was achieved by treating Col-0 plants with exogenous MeJA several days before water was withheld (**Supplemental Figure S4C**). The higher tolerance of JAR1-OE thus seems to be based on changes induced by the elevated JA-Ile content before the plants experienced drought stress. We observed that JAR1-OE plants were able to retain a relatively high water content of more than 80 % even after 18 days of water withdrawal (**Figure 2C**). This could be due to their smaller stomatal aperture and lower stomatal density observed even under control conditions (**Figure 7A and B**). Regulation of stomatal number and aperture diameter are important mechanisms to mitigate water deficiency. Exogenously applied MeJA was shown to regulate stomatal aperture in leaves (Raghavendra & Reddy, 1987; Hossain et al., 2011). The regulation of stomatal aperture, however, is a very complex process. The higher expression of *TGG1* and *TGG2* in JAR1-OE might play a role, since these myrosinases were shown to be involved in ABA- and MeJA-induced stomatal closure downstream of ROS production (Rhaman et al., 2020; Islam et al., 2009). The lower number of stomata in JAR1-OE already under normal growth conditions seem to be due to increased expression of genes that negatively regulate stomata patterning. This is consistent with the finding that exogenous MeJA application can negatively regulate stomatal development in cotyledons (Han et al., 2018).

Drought stress results in an accumulation of cytotoxic H_2_O_2_ and other ROS (Nocter et al., 2014). But ROS are also generated as secondary messengers and are involved in controlling hormone-dependent stress responses (Xia et al., 2015; Kwak et al., 2006). Controlled redox regulation is thus important to remove cytotoxic ROS levels, while sustaining ROS-dependent regulatory circuits. We could show that external addition of JA alleviates MV-induced H_2_O_2_ production in Col-0 but not in the *jar1-11* mutant. Previously, external MeJA application was reported to induce some genes involved in the ascorbate-glutathione cycle, one of the major mechanisms to adjust cytosolic H_2_O_2_ levels (Xiang & Oliver, 1998; Sasaki-Sekimoto et al., 2005; Zander et al., 2020). In our study, we observed upregulation of both *DHAR1* and *GR1/GR2* under drought in Col-0 but differential regulation of these genes in *jar1-11* and JAR1-OE (**Figure 7D and Supplemental Figure S6**). We did not see any difference in the expression of ascorbate-glutathione pathway genes under non-stress conditions in JAR1-OE, despite the increase in JA-Ile levels. Together, our data suggest that rather than generally inducing ascorbate-glutathione cycle activity, JA-Ile adjusts the flow through the ascorbate-glutathione cycle under drought conditions. Further studies are required to address this phenomenon, but it might play a role in making the JAR1-OE plants better able to deal with increased H_2_O_2_ levels upon drought.

### JA-Ile (and other jasmonates) play a role in drought stress priming

Jasmonates, especially JA-Ile, are at the core of JA-dependent stress responses through COI1-JAZ mediated transcriptional regulations. Interestingly, JAR1-OE plants not only showed an overall higher level of JA-Ile under control conditions but the net increase in JA-Ile under drought also exceeds that observed in Col-0 (**Figure 3**). Levels of the direct precursor JA increased in Col-0 upon drought, while they remain the same in JAR1-OE plants. This might indicate that under drought conditions, even the elevated levels of JAR1 proteins in Col-0 are not sufficient to convert all JA into JA-Ile. It is also likely that factors other than just the amount of JAR1 protein affect JA and JA-Ile levels under stress conditions. The basal level of JA-precursor *cis*-OPDA generally is almost 200 times higher compared to JA-Ile (de Ollas et al., 2015a; Balfagón et al., 2019; Figure 3) and under drought conditions, the magnitude of decrease in *cis*-OPDA content is much higher than the increase in JA-Ile. Together, this indicates that JA production from *cis*-OPDA is not the limiting factor for JA-Ile synthesis. However, the decrease in *cis*-OPDA is at a similar magnitude as the combined increase in JA, JA-Ile and their derivatives such a 12-OH-JA, 12-OH-JA-Ile and COOH-JA-Ile, all of which accumulate to a greater extent than JA-Ile itself. 12-OH-JA and 12-OH-JA-Ile were both found to modulate JA-Ile mediated gene expression, including genes involved in jasmonate biosynthesis (Miersch et al., 2008; Jimenez-Aleman et al., 2019; Poudel et al., 2019). They could thus play a role in balancing the jasmonate responses induced by JA-Ile. Their modulation of certain JA-Ile regulated processes seems to be based on the ability to also act as a bioactive ligands for the formation of COI1-JAZ receptor complexes (Poudel et al., 2019; Jimenez-Aleman et al., 2019). Thus, synthesis of JA-Ile together with the interconversion of JA and JA-Ile into various other derivatives might play an important part in stimuli-specific regulation but likely also in jasmonate homeostasis, i.e. by removal of excess JA-Ile in JAR1-OE under non-stress conditions. Especially intriguing in this respect is the high amount of the JA-derivative 12-*O*-JA-Glc that we found in Col-0, which is in contrast to published data from the Wasilewskija ecotype (Miersch et al., 2008) but is supported by recent findings in poplar (Ullah et al., 2019). Under drought stress conditions, its content increased in all lines, albeit to a lesser extent in JAR1-OE and a greater extent in *jar1-11*. 12-*O*-JA-Glc has been shown to accumulate 24 hours after wounding of tomato leaves and this accumulation was dependent on jasmonate biosynthesis (Miersch et al., 2008). It was suggested that it might be part of the pathway to remove JA-Ile accumulating under stress. While our study only shows the content of jasmonates at a single (and early) time point during the progressive drought stress, the data lead to the speculation of a stress induced continuous flow of JA-Ile synthesis and removal.

### Crosstalk between jasmonate and ABA signaling

Studies on various plant species have indicated that ABA and JA-signaling have a complex, interwoven relationship with regard to drought-stress priming and response. With regard to *MYC2* expression, it was shown previously that either jasmonate or ABA alone could induce its expression; however, the effect of both hormones together was much stronger (Lorenzo et al., 2004). This would explain the only slight increase of *MYC2* levels in JAR1-OE under control conditions, where ABA levels are not elevated. Liu and co-workers (Liu et al., 2016) proposed a model, in which exposure to drought activates transcription of *MYC2* via both ABA and jasmonate, which in the form of a positive feedback loop leads to further activation of JA synthesis and subsequently further elevated expression of jasmonate-dependent genes. This is in line with our finding that the expression of *MYC2*, as well as some genes involved in JA synthesis, is increased to a greater extent in JAR1-OE and to a lesser extent in *jar1-11* under drought conditions, when ABA levels are increased. Drought-induced ABA accumulation was evident in all three lines, but was enhanced in *jar1-11* and reduced in JAR1-OE compared to Col-0 (**Figure 3H**). Changes in ABA level corresponded to opposite alterations in the expression of genes related to ABA biosynthesis (**Figure 5C**). However, increase in expression of genes related to ABA biosynthesis in *jar-11* was accompanied by upregulation of genes involved in ABA degradation as well as of *ABI2*, a negative regulator of ABA signaling (Merlot et al., 2001), whose downregulation by exogenous MeJA application was recently described (Zander et al., 2020). Jasmonate signaling could thus also be part of a mechanism to reduce excessive amounts of ABA during drought stress conditions to keep the balance between drought protection and growth.

## Materials and methods

### Plant materials and growth conditions

All experiments in this study were performed on *Arabidopsis thaliana* (ecotype Columbia; Col-0) plants or transgenic lines created in the Col-0 background. The T-DNA insertion lines *jar1-11* (SALK_034543) and *jar1-12* (SALK_011510) were obtained from NASC (Nottingham Arabidopsis Stock Centre, UK) and plants homozygous for the T-DNA insertion were identified by PCR screening (a list of all primers used in this study can be found in **Supplemental Table S1**). Plants were grown either on ½ Murashige and Skoog medium (MS) medium (Duchefa Biochemie, Netherlands) with 1% sucrose and 0.6% [w/v] phytagel (Sigma-Aldrich, Inc., Germany) or on standard plant potting soil pretreated with Confidor WG 70 (Bayer Agrar, Germany). Plants grown on ½ MS were stratified for 2 days at 4^0^C in the dark. If not otherwise stated, plants were cultured in climatized growth chambers (equipped with Philips TLD 18W of alternating 830/840 light color temperature) at 22°C under long-day conditions (16 h light/8 h dark) with 100 µmol photons m^-2^ s^-1^. In some experiments, short-day conditions (8 h light/16 h dark) were applied.

### Generation of JAR1-YFP overexpression lines

To generate plants expressing JAR1.1 as a fusion protein with YFP under the control of the 35S promoter (35S::JAR1.1-YFP), the entire coding sequence of the *JAR1.1* variant was cloned into the pBIN19 vector (Datla et al., 1992) in frame with the *YFP* sequence using ApaI and NotI restriction sites. The resulting construct (**Supplemental Figure 1C**) was stably transformed into Arabidopsis wild type using the floral dip method. Three independent homozygous T-DNA insertion lines (JAR1-OE) were obtained each in the F3 generation. JAR1-YFP expression was confirmed through confocal microscopy, RT-qPCR, and western blot using an antibody against GFP (see below).

### Phenotyping and drought stress experiments

For phenotyping under normal and drought stress conditions, seeds were directly planted in potting soil. Five days later, young seedlings were transplanted to fresh pots containing 100 g potting soil (either one or four seedlings per pot). Plants were grown for 18 days with regular watering with identical volumes of tap water. Afterward, plants were either watered normally or were exposed to drought stress conditions by withholding watering for up to 14 days. During the drought-stress treatment, pot weights were measured regularly. The relative water content of soil (SWC) was calculated from the dried pot weight and adjusted between plant lines to ensure a similar drought stress level. SWC was calculated as {(pot weight at the time of stress)−(empty pot weight)}/{(initial pot weight)−(empty pot weight)} × 100. After SWC dropped to 10%, plants were rewatered with equal volumes of tap water and survival rates of plants were calculated 24 h and 7 d later. At least four independent experiments, each with several plants, were conducted for all experiments. The positioning of all pots in the climate chamber was randomized throughout the experiments. Photographs were taken at regular intervals and corresponding whole rosette leaves were collected for biochemical and RNA-seq analyses on day 32.

### Stomatal aperture, density and RWC measurements

Stomatal aperture diameters and density were measured from the 6th leaf of 21 day-old plants by collecting the leaf epidermis as described previously (Hossain et al., 2011). Briefly, excised rosette leaves were floated on a medium containing 5 mM KCl, 50 mM CaCl_2_, and 10 mM MES-Tris (pH 6.15) for 2 h in the light (80 µmol photons m^-2^ s^-1^). Subsequently, the abaxial side of the excised leaf was softly attached to a glass slide using a medical adhesive (stock no. 7730; Hollister), and then adaxial epidermis and mesophyll tissues were removed carefully with a razor blade to keep the intact lower epidermis on the slide. Pictures were taken immediately using the bright field option of a confocal microscope (SP8 Lightning, Leica, Weimar, Germany) and the aperture length was processed using the integrated LASX software.

The relative water content (RWC) of leaves was calculated according to (Barrs & Weatherley, 1962). Briefly, the weight of the whole rosette was measured immediately after collection (W) and after floating them on water for 24 h (TW). Finally, the dry weight (DW) was determined by fully desiccating the leaves in an oven at 70^0^C for 72 hours. RWC was then determined as [{(W-DW) / (TW-DW)**}** × 100].

### *In vivo* redox imaging

An Arabidopsis line carrying the cytosol-targeted roGFP2-Orp1 sensor previously described by Nietzel et al. (2019), was crossed with *jar1-11*, and homozygous plants of the F3 generation were used for imaging. *In vivo* redox imaging was performed on the leaves of 7-9 day-old seedlings as described in (Meyer et al., 2007) using a Leica SP8 lightning and data were processed using the integrated LASX software with the ‘quantify’ mode. In short, roGFP2 was excited at wavelengths 405 and 488 nm and the emission was detected from 505 to 530 nm. The ratiometric image of 405/488 nm was calculated based on a standardization using 50 mM DTT and 10 mM H_2_O_2_. Seedlings were pre-incubated in the imaging buffer (10 mM MES, 10 mM MgCl_2_, 10 mM CaCl_2_, 5 mM KCl, pH 5.8). Subsequently, seedlings were transferred into a perfusion chamber (QE-1, Warner instruments) to allow the exchange to different treatment solutions pumped through a peristaltic pump (PPS5, Multi-channel systems) under constant imaging. Pinhole was adjusted to 3. After each run, representative samples were calibrated with 10 mM DTT (ratio = 0.12) and 10 mM H_2_O_2_ (ratio = 3.0).

### Anthocyanin measurements

Anthocyanin content was measured according to the modified protocol of (Neff & Chory, 1998). Briefly, whole rosette leaves were ground in liquid N_2_, 100 mg of the ground tissue were mixed with extraction buffer (methanol with 1% HCl) and the mixture was placed at 4°C in the dark overnight. After the addition of 200 µl H_2_O and 500 µl chloroform, the samples were mixed thoroughly and centrifuged at 14,000 g for 5 minutes. After centrifugation, 400 µl of the supernatant was collected in a new tube and re-extracted with 400 µl of 60% Methanol 1% HCl, 40%. The absorbance of the solution was taken at 530 nm (anthocyanin) and 657 nm (background) and anthocyanin content was expressed as (A530-A657) per g fresh weight.

### Western blot analysis

For extraction of total proteins, leaf samples were first ground in liquid N_2_. Approximately 100 mg finely ground tissues were mixed with 100 µl 4 x SDS-PAGE solubilizing buffer, vortexed and then incubated at 96°C for 10 minutes. After centrifugation for 10 minutes at 14,000 g, proteins in the supernatant were separated on 10 % SDS-PAGE gels and blotted onto nitrocellulose membranes. Western blot analysis was performed by a standard protocol using an antibody against GFP (α-GFP, Roche) and a secondary antibody coupled with alkaline phosphatase (Pierce Goat Anti-Mouse, Thermofisher Scientific).

### Phytohormone analysis

Flash-frozen whole rosette leaves of 32 day-old Arabidopsis plants were ground to a fine powder in liquid N_2_. Approximately 50 mg of each sample was extracted with 1 ml methanol containing 3µl of a phytohormone standard mix (30 ng of D_6_-JA (HPC Standards GmbH, Cunnersdorf, Germany), 30 ng of D_6_-ABA (Santa Cruz Biotechnology, Dallas, TX, USA), 6 ng D_6_-JA-Ile (HPC Standards GmbH) as internal standards. The contents were vortexed vigorously for 4-5 seconds, incubated for 2 min at 25°C, and agitated at 1500 rpm in a heating block. The contents were then centrifuged at 13 000 xg at 4°C for 5 min. Approximately 900 µl of the supernatant was transferred to new 2 ml microcentrifuge tubes. The residual tissues were reextracted using 750 µl 100% methanol without standards. The supernatants (1650 µl in total) were completely dried under a flow of N_2_ at 30°C and redissolved in 300 µl 100% methanol.

Phytohormone analysis was performed on an Agilent 1260 high-performance liquid chromatography (HPLC) system (Agilent Technologies, Santa Clara, CA, USA) attached to an API 5000 tandem mass spectrometer (AB SCIEX, Darmstadt, Germany) as described by (Ullah et al., 2019). The parent ion and corresponding fragments of jasmonates and ABA were analyzed by multiple reaction monitoring as described earlier (Vadassery et al., 2012). The concentrations of ABA and jasmonates were determined as described previously by (Ullah et al., 2019).

### RNA extraction, cDNA synthesis and RT-qPCR

Total RNA was extracted from the whole rosette leaves of 32 day-old control and drought-stressed plants using the Quick-RNA Miniprep Kit (Zymo-Research, USA). RNA quality and quantity were determined using a Nabi UV/Vis Nano Spectrophotometer (LTF Labortechnik, Germany). For RT-qPCR analysis, cDNA was prepared from 1 µg of mRNA with RevertAid First Strand cDNA Synthesis Kit (Thermo Scientific, ThermoFischer Scientific). Gene expression was quantified using the Power SYBR Green PCR Master Mix (Applied Biosystems,

ThermoFisher Scientific) in 48 well-plates in a StepOne™ Real-Time PCR Thermocycler (Applied Biosystems, ThermoFisher Scientific) and the expression level was normalized to *Actin2* to express as relative quantity (2^-ΔΔCt^). A standard thermal profile was used with 50°C for 2 min, 95°C for 10 min, followed by 40 cycles of 95°C for 15 s and 60°C for 1 min. Amplicon dissociation curves were recorded after cycle 40 by heating from 60 to 95°C with a ramp speed of 0.05°C/s. Primers used for RT-qPCR are listed in Supplemental Table S1.

### RNA-seq analysis

For RNA-seq, the quality of RNA was checked by determining the RNA integrity number using a Tapestation 4200 (Agilent). The library preparation and sequencing were performed by the NGS Core Facilities at the University of Bonn, Germany. Approximately 200 ng of RNA was used for library construction. Sequencing libraries were prepared using the Lexogen‘s QuantSeq 3‘-mRNA-Seq Kit and sequenced on an Illumina HiSeq 2500 V4 platform with a read length of 1x50 bases. For each of the samples, three biological replicates were sequenced with an average sequencing depth of 10 million reads.

CLC Genomics Workbench (v.12.03, https://www.qiagenbioinformatics.com/) was used to process the raw sequencing data. Quality control and trimming were performed on FASTQ files of the samples. Quality trimming was performed based on a quality score limit of 0.05 and a maximum number of two ambiguities. To map the additional JAR1 reads from the JAR1.1-YFP lines, an additional chromosome comprising the YFP sequence was added to the Araport 11 (Cheng et al., 2017) genome and the annotation file. The FASTQ samples were then mapped to the modified Araport 11 genome, while only classifying reads as mapped which uniquely matched with ≥ 80% of their length and shared ≥ 90% identity with the reference genome. For the mapping to the gene models reads had to match with ≥ 90% of their length and share ≥ 90% similarity with a maximum of one hit allowed. Subsequently, counts for JAR1.1-YFP and JAR1 were combined. Further steps were completed using the R programming language (R Core Team 2020). To test the quality of the data, samples were clustered in a multidimensional scaling plot (MDS plot) using plotMDS. To assess differential expression of the sequencing data the Bioconductor package edgeR was used (Robinson et al., 2009). First the read counts were normalized by library sizes with the trimmed mean of M-values (TMM) method (Robinson & Oshlack, 2010). Then common and tagwise dispersion was calculated. For pairwise comparisons the exactTest function to calculate the p-value and the log2-fold-change were used. The resulting p-values were adjusted by using the False Discovery Rate (FDR) method (Benjamini & Hochberg, 1995). K-means clustering was performed using the kmeans function with the algorithm of Hartigan and Wong (1979) (Hartigan & Wong, 1979). The number of clusters for each clustering was estimated using the elbow method (Thorndike, 1953).

GO term enrichment analysis was performed with the topGO package (Alexa & Rahnenfuhrer, 2020). The athaliana_eg_gene dataset (Cheng et al., 2017) was downloaded from Ensembl Plants (Yates et al., 2020) via the BioMart package (Durinck et al., 2009). For this a weighted fisher test (Fisher, 1925) was run using the weighted01 algorithm (p ≤ 0.001). The resulting p-values were adjusted by using the BH method (Benjamini & Hochberg, 1995) filtering for an adjusted p-value ≤ 0.01.

Additionally, Transcripts Per Million (TPM) values were calculated based on the read counts. For individual genes, TPM values were compared by performing an ANOVA (Chambers et al., 1992) and a Tukey’s HSD test with a confidence interval of 0.95 (Tukey, 1949). Figures and plots were created using VennDiagram, pheatmap, ggpubr, and EnhancedVolcano included in the R package.

### Statistical analyses

Data were analyzed statistically with analysis of variance (ANOVA) followed by multiple comparisons (Tukey’s honest significant difference [HSD] test) in R v.4.0.3 using the ggplot2 package. One-way ANOVA was used to all parameters except hormonal data where two-way ANOVA was applied. For additional experiments, two-tailed t-test was used. Number of replicates and error bars are indicated in the figure legends. Bar plots with error bars were generated in Microsoft Excel, v.16.39. Real-time monitoring of the roGFP2-Orp1 sensor was done using the XY-simple linear regression with 95% confidence level in GraphPad Prismsoftware, v.9.0.0. Information on statistical processing for the RNA-seq are specified in the respective Methods section.

### Accession numbers

A list of accession numbers is provided in Supplemental Data Set S6.

## Supplemental Data

**Supplemental Data Set S1**: RNA-seq results from leaf tissue of 32 days old plants grown on soil.

**Supplemental Data Set S2**: List of DEGs found between control and drought conditions in each line and between Col-0 and *jar1-11* or JAR1-OE under control and drought conditions.

**Supplemental Data Set S3**: Selected DEGs sorted by biological processes.

**Supplemental Data Set S4**: GO term enrichment analysis

**Supplemental Data Set 5**: Hierarchical clustering of DEGs among wild-type, *jar1-11* and JAR1-OE.

**Supplemental Data Set S6**: List of Accession No.

**Supplemental Table S1:** List of Primers used in this study.

**Supplemental Figure S1:** Confirmation of the TDNA insertion into the *JAR1* locus and overexpression of JAR1.1-YFP.

**Supplemental Figure S2:** Effect of exogenous MeJA application on root growth.

**Supplemental Figure S3:** Phenotype of the *jar1-12* and additional JAR1-OE lines

**Supplemental Figure S4:** Drought stress phenotypes under long-day conditions

**Supplemental Figure S5:** Drought stress phenotypes under short-day conditions

**Supplemental Figure S6:** Proposed regulation of the ascorbate-glutathione cycle by JA-Ile.

## Acknowledgments

We are grateful to Prof. Dr. Frank Hochholdingher, INRES, Crop Functional Genomics, University of Bonn for facilitating the RNA-seq data analysis in his group. We acknowledge the NGS Core Facility, University of Bonn, for providing the RNA-seq service. Finally, we thank Dr. Fatima Chigri for careful proofreading of the manuscript.

## Author Contributions

S.M. and U.C.V. designed the research. S.M., C.U., and S.B. performed the research. S.M., C.U., A.K., P.Y., J.G., and U.C.V. analyzed the data. S.M., and U.C.V. wrote the paper with input from all authors.

